# Comparative Transcriptomics of *Arabidopsis*, *Medicago*, *Brachypodium* and *Setaria* species during Phosphorus limitation

**DOI:** 10.1101/2024.07.02.601706

**Authors:** Pooja Pant, Hui Duan, Nick Krom, Wolf-Rűdiger Scheible

## Abstract

Translating biological knowledge from *Arabidopsis* to crop species is important to advance agriculture and secure food production in the face of dwindling fertilizer resources, biotic and abiotic stresses. However, it is often not trivial to identify functional homologs (orthologs) of *Arabidopsis* genes in crops. Combining sequence and expression data can improve the correct prediction of orthologs. Here, we conducted a large-scale RNA sequencing based transcriptomics study of *Arabidopsis*, *Medicago*, *Brachypodium* and *Setaria* grown side-by-side in Phosphorus (P)-sufficient and P-limited conditions to generate comparable transcriptomics datasets. Comparison of top 200 P-limitation induced genes in *Arabidopsis* revealed that ∼80% of these genes have identifiable close homologs in the other three species but only ∼50% retain their P-limitation response in the legume and grasses. Most of the hallmark genes of the P-starvation response were found conserved in all four species. This study reveals many known, novel, unannotated, conserved and species-specific regulations of the transcriptional P-starvation response. Identification and experimental verification of expressologs by independent RT-qPCR for P-limitation marker genes in *Prunus* showed the usefulness of comparative transcriptomics in pinpointing the functional orthologs in diverse crop species. This study provides an unprecedented resource for functional genomics and translational research to create P-efficient crops.

**HIGHLIGHT:** Comparative transcriptomics reveals novel, known, conserved and specific transcriptome responding to Phosphorus limitation in *Arabidopsis, Medicago, Brachypodium* and *Setaria* to facilitate translational research in crops.

## INTRODUCTION

Plant-available phosphorus (P) is often in short supply in agricultural soils and its low availability limits crop production. P is a structural component of energy metabolites, membrane lipids and nucleic acids, and is involved in signal transduction cascades. Protein phosphorylation acts as a precise molecular switch to turn on and off the specific signaling pathways needed for plant growth or defense in rapidly changing environments. (Zhang et al., 2023). Large amounts of mineral P fertilizers are applied to soil to meet the crops’ P demand (Marschner, 1995; Raghothama, 1999; Poirier and Bucher, 2002). Application of P fertilizer is both costly and ecologically hazardous as the runoff from agricultural field causes eutrophication of aquatic bodies. Minable global P resources are geographically restricted, non-renewable, and finite. Therefore, research on plant responses and their regulation during P-limitation, and translation of the obtained knowledge to improve P-use efficiency in agricultural production systems holds economic, political, and environmental impetus.

P-limitation leads to considerable changes in gene expression in plants. A number of studies described changes in gene expression during P-limitation in the model plant *Arabidopsis*, which include studies with a limited microarray (Wu et al., 2003), ATH1 GeneChip (Misson et al., 2005; Morcuende et al., 2007; Bustos et al., 2010; Lin et al., 2011; Hoehenwarter et al., 2016), tiling array (Woo et al., 2012), Quantitative reverse transcription polymerase chain reaction (RT-qPCR) (Morcuende et al., 2007; Pant et al., 2009) and small RNA sequencing (Hsieh et al., 2009; Pant et al., 2009). Transcriptome studies during P-limitation have also been carried out in other species including rice (Wasaki et al., 2006; Secco et al., 2013; Tyagi and Rai, 2016), wheat (Oono et al., 2013), maize (Calderon-Vazquez et al., 2008; Lin et al., 2013; Pei et al., 2013; Du et al., 2016; Sun et al., 2016), Masson pine (Fan et al., 2014), Harsh Hakea (Kuppusamy et al., 2014), bean (Hernandez et al., 2007), white lupin (Secco et al., 2014) or soybean (Wang et al., 2016; Zhang et al., 2016). Despite such efforts, there exists considerable variation in terms of the number of P-limitation responsive genes and extent of their responses among studies. Experimental variations and parameters certainly account for some differences between studies. Therefore, it is also important to grow the plants together under the same experimental setting for better comparison and identification of orthologues. A comparative transcriptome analysis of P-responsive genes in shoots and roots of different plant species by growing them side-by-side in same conditions has not been reported.

Homologs are the genes related by common ancestry that can be subcategorized into two main categories: orthologs and paralogs (Fitch, 1970; Koonin, 2005; Glover et al., 2016). Orthologs are the genes that evolved from a common ancestral gene by speciation in different species. Normally, orthologs retain the same function, while paralogs, which are the genes related by genomic duplication, are supposed to evolve new functions (Tatusov et al., 1997). Identification of orthologs is a key tool for predicting gene function in new species as well as for meaningful comparison of gene and genome organization of different species (Fitch, 1995; Tatusov et al., 1997). There exists one-to-one, one-to-many or many-to-many orthologous relationships between the genes in different species due to a series of speciation and duplication events of an ancestral gene (Tatusov et al., 1997). To identify the homologs in different species, one has to rely on sequence similarity and suitable statistics (Moreno-Hagelsieb and Latimer, 2008). Precise ortholog detection in different species is hampered by several evolutionary events including gene duplication, loss, fusion/fission, and segmental and whole-genome duplication (Tekaia, 2016) as well as insertion, deletion or substitution. Combining sequence and expression data has been shown to improve the correct prediction of functional orthologs across the different species (Bergmann et al., 2004; Das et al., 2016).

Ornamental, flowering cherry trees (*Prunus* species) are popular all over the world. Flowering cherries are the second most valuable deciduous flowering trees in the United States (Guo et al., 2018). P-limitation is known to cause retarded plant growth and development, earlier abscission, inferior flower quality, decreased number of flowers and low flowering intensity (Erel et al., 2008; Marschner and Marschner, 2012; Erel et al., 2016). P is an essential element for optimal flower production in *Prunus* species. However, there is no data to understand the P-homeostasis mechanism in *Prunus* species and how similar or different it is compared to other plant species.

As the first and most widely used organism in the plant science research community, *Arabidopsis* continues to be the best-studied model plant species and offers a wealth of information including plant responses to P-limitation, P-limitation responsive transcriptome, and signaling/regulatory pathways. However, it is largely unknown whether the P-responsive genes and the extent of their response to P-limitation observed in *Arabidopsis* is conserved in model legumes and grasses. RNA sequencing (RNA-seq) based high-throughput sequencing can provide information about the genome-wide transcriptome in a biological sample and is better than microarrays in detecting lowly expressed genes (Wang et al., 2014). To this end, this study has taken into consideration a large-scale RNA-seq based transcriptome profiling of model plants for agriculturally important species like *Medicago truncatula* (model for legumes), *Brachypodium distachyon* (C_3_ grass, similar to rice or wheat) and *Setaria viridis* (C_4_ grass, similar to switchgrass or maize) grown under P-limitation condition and compare the P-limitation transcriptome responses to *Arabidopsis thaliana*. Growing four different plants species side-by-side in the same condition facilitated the direct and more reliable comparison of the orthologues and overall transcriptome responding to P-limitation. Genome-wide comparison of the phosphate starvation response in four plant species showed that the general Pi-starvation response is similar across different plant species, but there are also many species-specific differences. This study also established experimental protocol to study phosphate/nutrient homeostasis in *Prunus*, identified P-limitation marker genes and verified induction of these genes experimentally using reverse transcription-quantitative polymerase chain reaction (RT-qPCR). The importance of comparative transcriptomics in translating knowledge from model species to other evolutionarily close and distant species is also discussed.

## MATERIALS AND METHODS

### Plant materials and growth condition

*Medicago truncatula* subspecies Jemalong A17 (HM101) (http://www.Medicagohapmap.org) seeds were obtained from Dr. Nevin Young (University of Minnesota). *Arabidopsis thaliana* ecotype Col ‘0’ seeds were in the group resource, originally obtained from The *Arabidopsis* Information Resource (TAIR; https://www.arabidopsis.org/); *Brachypodium distachyon* line Bd21-3 and *Setaria viridis* A10 seeds were obtained from Dr. Zengyu Wang’s lab at The Samuel Roberts Noble Foundation. All seeds were multiplied in the standard greenhouse condition at The Samuel Roberts Noble Foundation before use in molecular and phenotypic experiments. Same batch of the seeds were surface sterilized and kept for synchronization at 4°C for 3 days. *Medicago*, *Brachypodium* and *Setaria* seedlings were germinated in dark and kept under white light for two days before transplanting to pots containing sand and perlite mixture (1:3). *Arabidopsis* seeds were germinated and grown as described previously (Scheible et al., 2023). All the plants were divided into two groups and were watered with the full nutrient solution with (FN or +P) and without phosphate (-P) for the first three days and then the nutrient solution was diluted 5X and supplied to the plants every other day. Nutrient media was prepared as described previously (Morcuende et al., 2007; Scheible et al., 2023). Nineteen-day-old plants showed typical P-limitation phenotype (dark-green leaves and smaller plants) and confirmed to have >10-fold reduction in inorganic phosphate (Pi) content in the leaves in all P-limited plants in preliminary experiments. Measurement of Pi-content and induction of known *Arabidopsis* P-limitation marker genes and their putative orthologs tested in other three species in preliminary experiments of this study was used to determine the best age of the plants to harvest shoots and roots for RNA-seq experiments. The roots and shoots were harvested, flash frozen in liquid Nitrogen and stored at -80°C until further use. Reduction in Pi-content and induction of selected putative marker genes in -P samples as compared to +P samples were confirmed by RT-qPCR before RNA-seq library preparation and sequencing. Three independently prepared batches of plants grown under +P and -P conditions were used for RNA-seq experiments. Materials from five individual plants were pooled for each of the root and shoot samples as described previously (Scheible et al., 2023).

### Phosphate (Pi) measurement, RNA isolation, cDNA synthesis, and RT-qPCR

Measurement of the inorganic phosphate was done using the colorimetric micromethod (Itaya and Ui, 1966) as described in (Pant et al., 2015; Scheible et al., 2023). Briefly, frozen plant material was ground in liquid nitrogen, weighed (25 mg) and soluble inorganic phosphate was extracted in 250 µl Milli-Q water (Millipore Sigma) by repeatedly (3X) freezing and heating the samples at 95°C for 5 minutes. After centrifugation at 13,000 rpm for 15 minutes, 10 µl of the 5 times diluted supernatant was mixed with 100 µl HCl and 100 µl malachite green color reagent [1 vol. 4.2% (NH_4_)_6_ Mo_7_O_24_.H_2_O in 5 M HCl and 3 volumes of 0.2 % Malachite Green dye in water] in a 96 well flat-bottomed microtiter plate. The plate was incubated at room temperature in dark for 15 minutes and the absorbance was measured at 630nm using a Plate reader Infinite 200 PRO. RNA isolation, DNase treatment, cDNA synthesis, and RT-qPCR gene expression analysis was performed with three independent biological replicates and two technical replicates as described previously (Scheible et al., 2023). NanoDrop Spectrophotometer (ND-8000) was used to check the concentration and purity of RNA, and Agilent Bioanalyzer 2100 was used to check the RNA integrity. RNA samples with RNA Integrity Number (RIN) values >8 were taken for RNA-seq.

### RNA-seq deep sequencing and read processing

RNA sequencing and read processing was performed as described previously (Scheible et al., 2023). Briefly, RNA-seq libraries were prepared using the Illumina® TruSeq® RNA Sample Preparation Kit, according to manufacturer’s instructions. Validation of the RNA-seq libraries was performed by RT-qPCR before loading and sequencing on a flow cell of Illumina HiSeq2000 sequencing instrument at BGI Americas (www.bgiamericas.com). RNA-seq data from *Arabidopsis thaliana*, *Brachypodium distachyon*, *Medicago truncatula* were mapped to their respective genomes (TAIR10 for *Arabidopsis*, Phytozome v2.1 for *Brachypodium*, and JCVI v4.0 for *Medicago*) and RNA-seq reads from *Setaria viridis* was mapped to the *Setaria italica* (Phytozome v2.1) genome which showed ∼80% similarity as well as to its own *Setaria viridis* genome (Phytozome v2.1). RNA-seq data was analyzed using HISAT and StringTie (Kim et al., 2013; Pertea et al., 2016) as described previously (Scheible et al., 2023). Fold changes were calculated by adding 0.1 to each Fragments Per Kilobase of transcript per Million mapped reads (FPKM) value, to avoid divisions by zero. To identify the P status-responsive gene transcripts (PRGTs), the following filtering criteria were used: shoot PRGTs had to display a ≥5-fold average change for the two biological replicates, with each replicate being at least 3-fold changed (average [R1+R2]/2 ≥5; R1 ≥3 and R2 ≥3 or average [R1+R2]/2 ≤0.2; R1<0.333 and R2≤0.333). Root PRGTs were selected if the average change for two replicates was at least 5-fold with each of the two replicates showing an at least 3-fold change, and the third replicate showing at least a 2-fold change (average [R1+R2]/2 ≥5; R1 ≥3 and R2 ≥3 and R3 ≥2 or average [R1+R2]/2 ≤0.2; R1<0.333 and R2 ≤0.333 and R3 ≤0.5). The existence and P status response of the known and unannotated novel genes was confirmed by visualizing the read alignment (Binary Alignment/Map, BAM files) in integrative genomics viewer (IGV version 2.3.72; https://software.broadinstitute.org/software/igv) (Thorvaldsdottir et al., 2013) and performing RT–qPCR expression analysis for some selected genes/loci (Scheible et al., 2023).

### Gene ortholog identification, phylogenetic trees and websites used

Sequence information and phylogenetic analysis was used to identify the putative orthologous genes. Protein and nucleotide sequences were obtained from Phytozome database (http://www.phytozome.net) (Goodstein et al., 2012), TAIR10 www.arabidopsis.org (Berardini et al., 2015), LegumeIP (Li et al., 2016) and EnsemblPlants database (http://plants.ensembl.org/index.html,) (Kersey et al., 2016). The EnsemblPlants database and comparative genomics platform in PLAZA3.0 (http://bioinformatics.psb.ugent.be/plaza/) (Proost et al., 2015) was used for ortholog identification in different species. Some of the orthologous genes were identified and/or re-confirmed by one-by-one reciprocal best hit (RBH) BLAST analysis. Phylogenetic trees and the evolutionary history was inferred by using the Maximum Likelihood method in MEGA software (Tamura et al., 2013) and Geneious7.1.8 (http://www.geneious.com), (Kearse et al., 2012). Protein domain analysis was performed using InterProScan (https://www.ebi.ac.uk/interpro/search/sequence-search), UniProt (http://www.uniprot.org/) (UniProt, 2015) and conserved domain database (CDD) (http://www.ncbi.nlm.nih.gov/Structure/cdd/wrpsb.cgi) (Marchler-Bauer, et al. 2015). The nucleotide sequences were translated to protein sequence using the ExPASy-Translate tool (http://web.expasy.org/translate/). Gene expression overview for specific genes as well as information about homologous genes was checked in eFP Browser of MedicMine at http://MedicMine.jcvi.org/ (Smith et al., 2012) and *Arabidopsis* eFP browser at http://bar.utoronto.ca/efp/cgi-bin/efpWeb.cgi. MapMan ontologies were retrieved from MapMan database at http://mapman.gabipd.org (Thimm et al., 2004; Usadel et al., 2005). Gene families were identified using InterPro database (www.ebi.ac.uk/interpro).

### Establishment of in vitro culture system for P-limitation study in *Prunus*

*Prunus* tissue culture plants were originally derived from shoot apical meristems (SAM) of a *Prunus* hybrid, (*Prunus maackii* X *Prunus campanulate*). Shoots were *in vitro* propagated in MS basal medium with vitamins (M519) supplemented with 30 g/L sucrose (Sigma, S5391), 4.5 µM (1 mg/L) 6-benzylaminopurine (BA) and 0.5 µM (0.1 mg/L) 1-Naphthaleneacetic acid (NAA). The medium was adjusted to pH 5.8 using 1N KOH prior to addition of 6 g/L Phytagar (Gibco, Life Technologies, U.S.A) and autoclaving for 20 min at 121°C. *In vitro* shoots were sub-cultured to a fresh medium in sterile PhytoCon vessels (C215) every six weeks. Plants were maintained for more than six months in a Percival growth chamber. For P-limitation experiments, uniform explants (SAM) were taken from the mass of shoots proliferated from the same mother explant and freshly cultured on MS agar media without hormones. Shoots were rooted and six-weeks-old uniform, rooted plants were selected, rinsed with sterile deionized water twice before transferring in -P and +P media in Majenta jars. For preparing -P and +P media, MS modified basal salt mixture without NH_4_NO_3_, KNO_3_, or KH_2_PO_4_ (M407) was supplemented with 20.6 mM NH_4_NO_3,_ 18.8 mM KNO_3,_ 200 µM MES, 3 mM KH2PO4 (+P) and 2.5 mM KCl (-P) before adjusting to pH 5.8 and adding 6 g/L Phytagar. Chemicals were obtained from PhytoTechnology Laboratory, KS, USA, unless otherwise noted. The media was autoclaved and 150 ml was poured into each Magenta vessel (Sigma, V8380). A polypropylene coupler (Sigma, C0667) was used for joining two Magenta vessels and sealed with 1inch Micropore (3M) medical gas-permeable tape to allow for air exchange. Plants were grown under -P and +P condition in Magenta jars in a Percival growth chamber for 25 days until the typical P-limitation phenotype (dark-green leaves and growth retardation) was obvious. Plant positions within the growth chamber were randomized every two days to avoid position effects. For harvest, plants were quickly washed with deionized water, shoots and roots separated and immediately flash-frozen in liquid N_2_ before storage at -80°C until further analysis.

## RESULTS

### RNA-seq study identified many known and novel P limitation-responsive gene transcripts in *Arabidopsis*, *Medicago*, *Brachypodium* and *Setaria*

*Arabidopsis*, *Medicago*, *Brachypodium* and *Setaria* plants grown side-by-side in pots under P-sufficient (+P) and P-limitation condition generated a robust comparable transcriptomics dataset. Plants from all four species grown under P-limited conditions showed typical symptoms of P-starvation including reduced growth, dark green leaves, and accumulation of anthocyanin pigment (Figure 1A). Pi measurements showed that P-sufficient plants had ∼3 mg /g FW of Pi, whereas it was strongly (>10-fold) reduced in all P-limited plants (Figure 1B), confirming the visual P-starvation. P-limitation at the molecular level was confirmed by RT-qPCR expression analysis of P-starvation-inducible marker genes (Figure 1C). *Arabidopsis NON-SPECIFIC PHOSPHOLIPASE C* (*NPC5, AT3G03540*), *TRANSCRIPTION ELONGATION FACTOR* (*SPT5, At2g34210*) and *INDUCED BY PHOSPHATE STARVATION1* (*IPS1*, *At3g09922*) showed strong induction in P-limited *Arabidopsis* plants. Since the P-limitation marker genes for *Medicago*, *Brachypodium* and *Setaria* were not known, marker genes were selected in these species based on the sequence homology to known *Arabidopsis* P-limitation marker genes (Bari et al., 2006; Morcuende et al., 2007). The selected P-limitation marker genes, *SPX* (*<u>S</u>YG1<u>/P</u>ho81<u>/X</u>PR1*) domain gene (*Medtr1g012440*), *miR399h* and *miR399i* in *Medicago*, *SPX* (*LOC100840599*), *miR399a* and *miR399b* in *Brachypodium*, and *SPX* (*LOC101762844*), *NON-SPECIFIC PHOSPHOLIPASE C* (*PPl, Sevir.9G027300*) and *GLYCEROPHOSPHORYL DIESTER PHOSPHODIESTERASE* (*GDPD, Sevir.9G171400*) in *Setaria*, showed strong induction (>25-fold) in P-limited plants (Figure 1C).

**Figure 1.**
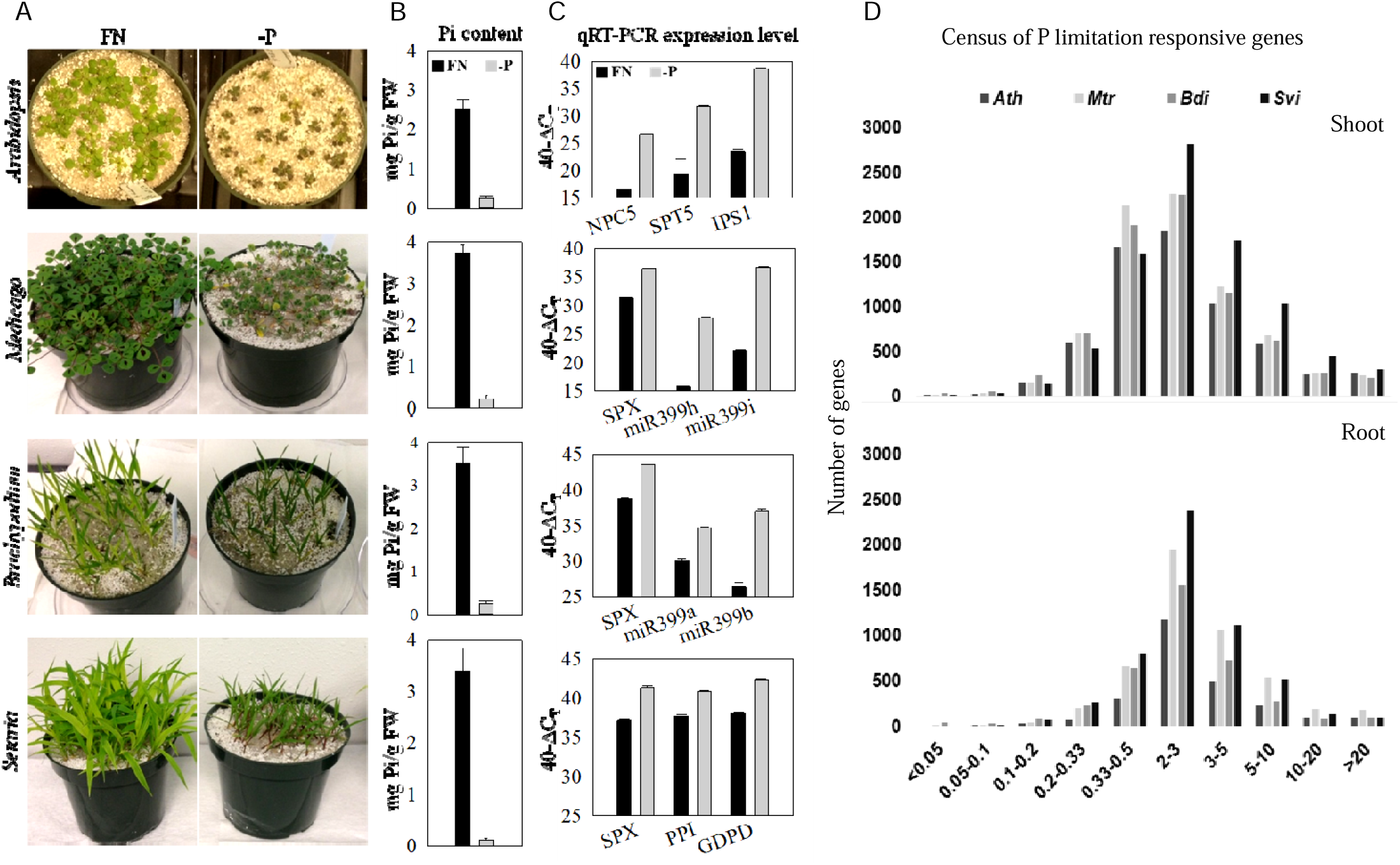
Phenology of experimental plants and differential transcriptome overview. (A) *Arabidopsis thaliana*, *Medicago truncatula*, *Brachypodium distachyon* and *Setaria viridis* plants were grown side-by-side in full nutrition (FN, left photos) and P-limitation (-P) (right photos) condition. (B) The concentration of inorganic phosphate (Pi mg/g FW, average ± standard deviation, *n*=5) for each of the four species (black bars = FN and white bars = P-limitation). (C) Identification and quantification of P-limitation marker gene expression using RT-qPCR in the shoots of the four species. Transcript abundance is plotted on a log_2_ scale as 40-ΔC_T_ with ΔC_T_ being the difference between the C_T_ (threshold cycle number) of the gene of interest and the invariant reference gene. ΔC_T_ of 5 equals ∼32-fold difference in expression. (D) Census of P-limitation responsive genes. The fold change range of differentially expressed genes in each of the four plant species is shown along the x-axis and the number of genes in each fold change range/category are shown along the y-axis. Fold change (-P *vs.* FN) was calculated from the RNA-seq expression data for shoot and root of *Arabidopsis (Ath)*, *Medicago (Mtr)*, *Brachypodium (Bdi)* and *Setaria (Svi)*.

The output file from the RNA-seq and HiSat StringTie analysis pipeline contains 26,922 gene entries of which 25,927 have an assigned reference in *Arabidopsis*; 38,380 gene entries with 34,994 assigned references in *Medicago*; 31,398 gene entries with 27,900 assigned references in *Brachypodium*; and 34,255 gene entries with 30,301 assigned references in *Setaria* (Table S1). Analysis of the RNA-seq data and identification of PRGTs was performed with strict criteria as described in the methods section. We identified a total of 2,347 PRGTs in *A.th.*, 3,562 in *B.di.*, 3,350 in *M.tr.* and 2,977 in *S.vi.* that were differentially expressed with ≥3-fold change in shoots and in roots or both shoots and roots respectively (Figure 1D). Of these, 229, 614, 484 and 626 PRGTs did not have assigned references in *A.th.*, *B.di.*, *M.tr.* and *S.vi.*, respectively (Table S1, Figure 2A and 3B). Among these PRGTs, there were 610, 920, 887 and 595 genes in shoot and 322, 427, 515 and 418 genes in roots that were highly induced (≥5-fold) and 180, 323, 201 and 184 genes in shoot and 59, 240, 134 and 86 genes in root that were highly repressed (≥5-fold) in *A.th.*, *B.di.*, *M.tr.* and *S.vi.* respectively (Figure 2A).

**Figure 2.**
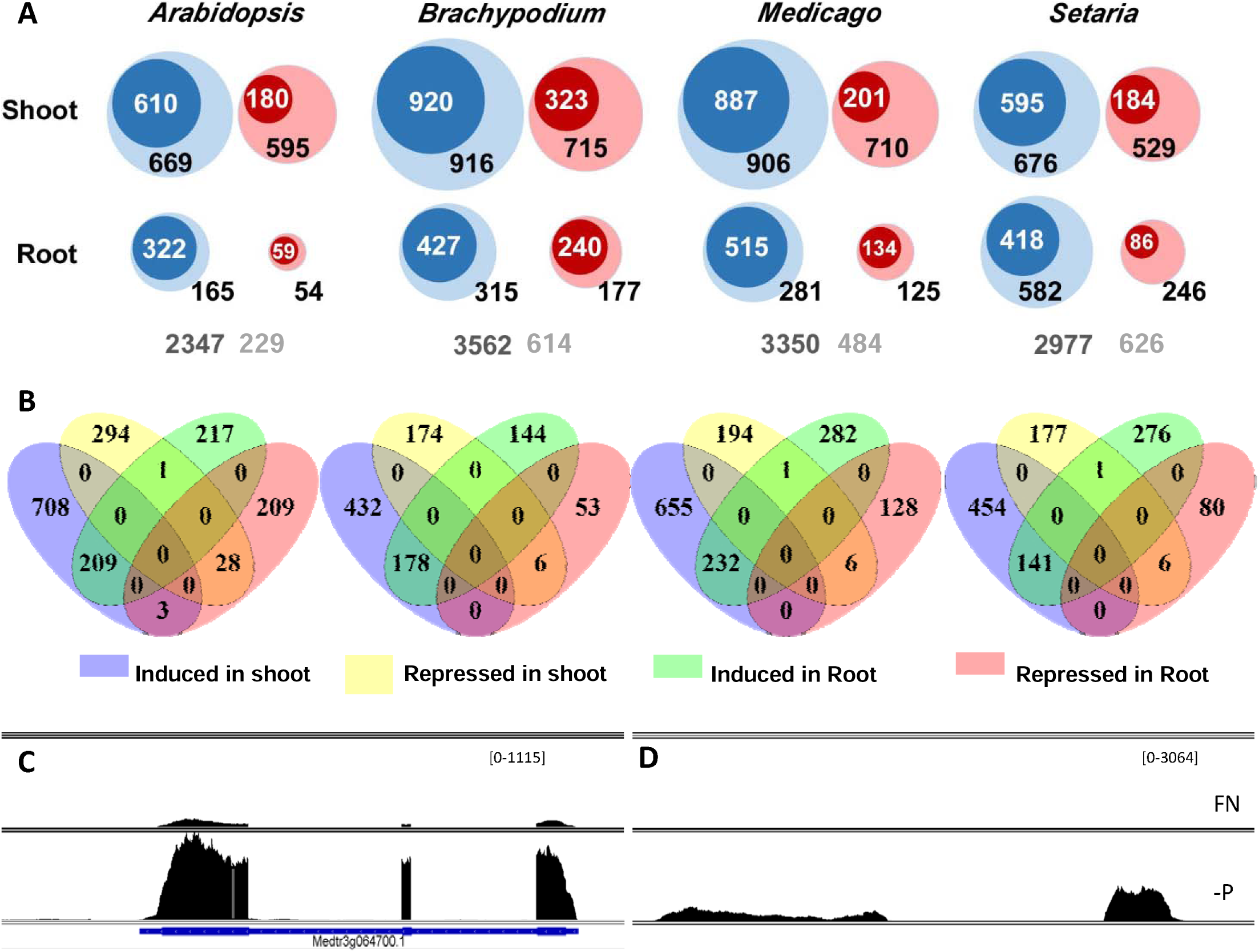
P status-responsive gene transcripts (PRGTs) identified by RNA-seq in shoots and roots of *Arabidopsis*, *Medicago*, *Brachypodium* and *Setaria* are shown. (A) Blue circles indicate the numbers of induced and red circles indicate repressed gene transcripts in the shoot (top row) or root (bottom row). Circle size is proportional to gene number. Numbers in white font indicate ≥5-fold changed, and numbers in black font indicate the number of gene transcripts that are ≥3-fold and <5-fold changed. The numbers in dark gray font indicate total number of PRGTs with ≥3-fold change in shoots and in roots or both shoots and roots and light grey font indicate those without any assigned references (novel genes). (B) Venn diagram of PRGTs (≥3-fold changed) showing the overlap of the numbers of genes induced or repressed in shoots or roots in four plant species. Fold change was calculated as described in methods. (C and D) Sashimi plots of P-limitation induced known and novel genes in Medicago. (C) *Medtr3g064700* annotated gene and (D) an unannotated (novel) gene in the Medicago visualized using IGV and BAM-files originating from RNA sequencing of shoots of full nutrition (FN, upper track) and P-limited (-P, lower track). Scaling of FPKM read numbers is the same for plots of the same transcript as indicated in parentheses (upper right corner of each panel). In each panel, X-axes indicate genomic locations and Y-axis indicate transcription intensity in shoot samples.

**Figure 3.**
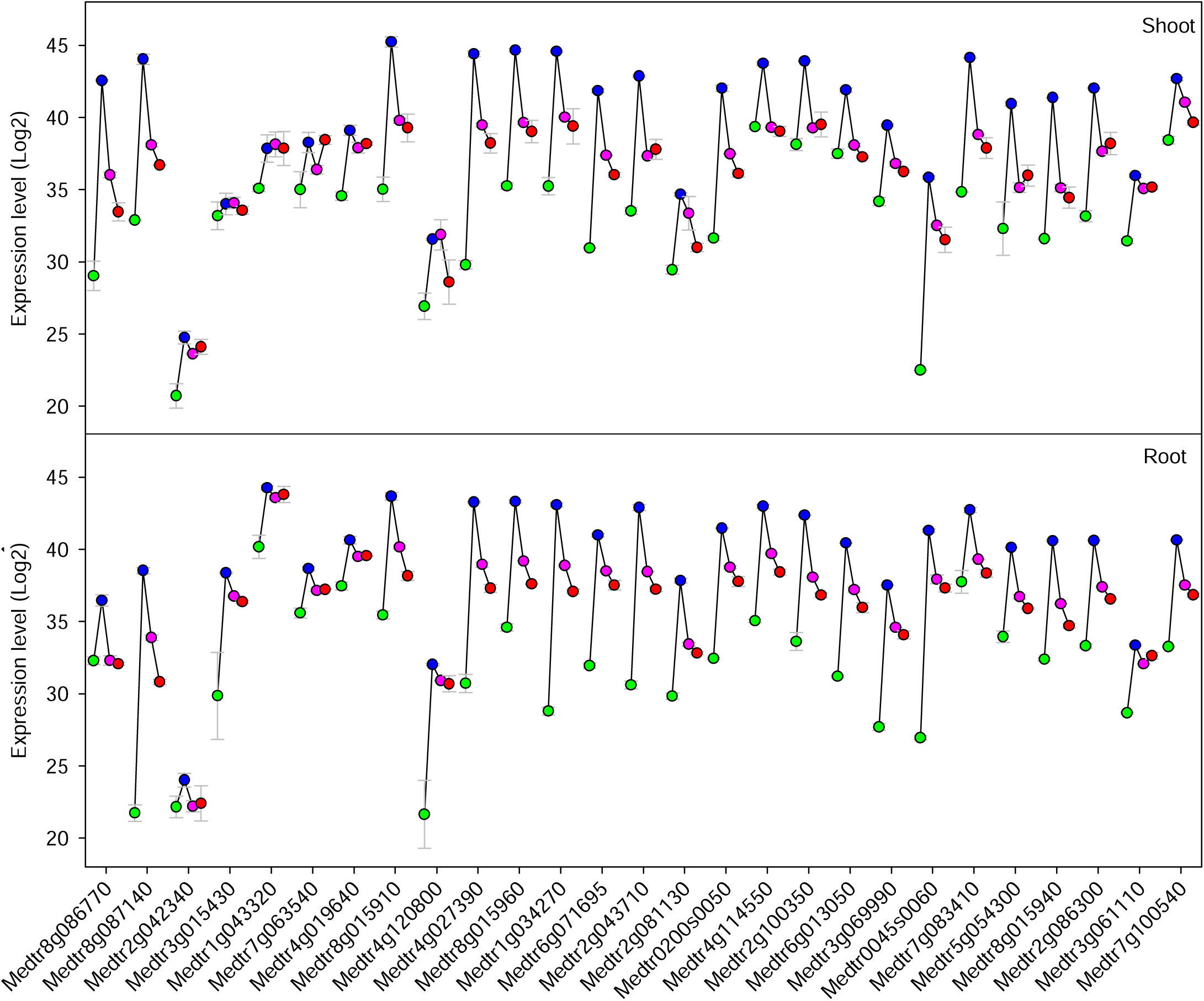
Induction of PSI genes reversed after P-resupply to P-starved plants and novel PSI genes. Proof of concept for randomly selected twenty-eight *Medicago* genes identified as phosphate starvation-inducible (PSI) by RNA-seq were confirmed for their reversible induction by P-limitation using RT-qPCR analysis in shoot and root samples. The RT-qPCR analysis was performed on the plants grown under full nutrition (FN), P-limitation (-P), and after 3 hours (P3hr) and 6 hours (P6hr) after P resupply to the P-starved plants. Expression level for each gene is given on a double logarithmic scale as 40-ΔC_T_, where ΔC_T_ is the difference between the threshold cycle number (C_T_) for each gene and the reference gene (*Ubiquitin*) and the error bars represent Standard deviation of two biological replicates. The experiment was repeated three times with similar results. Since most of these genes are not annotated/characterized yet (e.g. hypothetical proteins), gene IDs are shown along the X-axis for consistency.

We noticed that some genes responded in an opposing manner in shoot and roots, being induced in shoots but repressed in roots, and *vice versa* (Figure 2B). The opposing response of these genes could be due to their tissue-specific regulation, alternative splicing (as some splice forms are found in shoot and others are found in root), or epigenetics (Widman et al., 2014) that would help them to perform different, tissue- or organ-specific roles in plants to maintain P-homeostasis. Many P-starvation inducible (PSI) genes were expressed at a very low level during normal (P-sufficient) condition (Figure 2C), while some were not expressed at all during normal condition but were highly induced during P-limitation (Figure 2D). Several new *PSI* genes with reference and without reference (unannotated in the genome) were identified by RNA-seq analysis (Table S1, Figure 2). Most of these novel genes were expressed only during P-limitation with very little or no expression during P-sufficient conditions (Figure 2D). Careful analysis of the gene expression using IGV visualization of the RNA-seq reads revealed that many genes undergo alternative splicing during P-limitation in all four species (Figure S1). Well known P starvation inducible genes families such as purple acid phosphatases, phosphate transporters, transcription factors, methyl transferases, SPX-domain containing and PHO1-like genes, kinases, ribonucleases, protein inhibitors, endonucleases, glycolipid synthesis, senescence and pathogenesis-related genes, hypothetical proteins, or uncharacterized genes were among the top 100 PSI genes in all four species. These genes further confirm that the plants of the different species experienced similar P-limitation.

### Induction of P-starvation induced genes attenuated after P-resupply to P-starved plants

To verify the results obtained from the RNA-seq analysis, a subset of the highly phosphate starvation inducible (PSI) genes in shoots and/or roots were checked for their P starvation responses by RT-qPCR. Independently grown plant materials in P-sufficient (+P), P-limited (-P), and after 3 hr and 6 hr of Pi resupply to P-starved plants were used for verification of these *PSI* genes. RT-qPCR analysis revealed that almost all the selected genes were induced by P-limitation and their expression was dropped after P resupply to the P-starved plants, which confirms their expression is regulated by P availability. These highly *PSI* genes may have important roles in P homeostasis in plants and can be selected as marker genes for future P-limitation studies in *Medicago* (Figure 3) and *Brachypodium* (Figure S2). Among these highly P-limitation induced genes in *Brachypodium* are the genes involved in glycerophosphodiester phosphodiesterase activity, phosphatase activity, inorganic phosphate transmembrane transporter activity, triacylglycerol lipase activity, sugar:hydrogen symporter activity, and organophosphate:inorganic phosphate antiporter activity (Blast2GO). The exact function of most of these highly *PSI* genes in *Brachypodium* are not known yet. In *Medicago*, the strong phosphate starvation-inducible genes (Figure 3) include those encoding hypothetical or transmembrane proteins, 2,3-diketo-5-methylthio-1-phosphopentane phosphatase, putative phosphoglucomutase, phosphorus starvation-induced protein, purple acid phosphatase family proteins, IDS4-like proteins (SPX), inorganic pyrophosphatase, purple acid phosphatase, cytochrome P450 family 90 protein, glycerophosphoryl diester phosphodiesterase family protein, myb-like transcription factor family protein, alpha/beta hydrolase family protein, subtilisin-like serine protease and thioredoxin-like protein. Reversible expression of these gene transcripts suggests that their expression during P-limitation is a direct response to P limitation rather than an indirect response to altered growth/development that occur due to P limitation.

### Phosphate starvation-induced hallmark gene families are largely conserved

We analyzed the presence and response of well-known P starvation-inducible hallmark gene transcripts in *A.th.*, *M.tr.*, *B.di.* and *S.vi.* These include transcripts encoding purple acid phosphatases (PAP), phosphate transporters (PHT), SPX-domain and PHO1-like proteins, ribonucleases (RNS), glycolipid synthases and lipases, endonucleases, and phosphoenolpyruvate carboxylase kinase (PPCK) (Figure 4). Many of these derive from multigene families. We found that the responses of these transcripts are conserved across all four species, but the family size and number of family members showing a strong response (mainly induction) to P-limitation vary across species. There are 29, 38, 29 and 39 members in the *PAP* gene family in *A.th.*, *M.tr.*, *B.di.* and *S.vi.*, and out of these 27, 29, 22 and 31 were detected by RNA-seq, and 14, 13, 13, and 10 genes, respectively, were induced by ≥2-fold in shoot and/or root. Several transcriptional regulator (TR) genes were also regulated differently during P-limitation in all four plant species (Figure S3). responding to phosphorus limitation in four plant P-responsive hallmark gene families such as SPX, PHTs and glycolipid synthesis genes were investigated in more detail by constructing phylogenetic trees and comparing the P-response in individual subclades in different species (Figure 5, S4 and S5).

**Figure 4.**
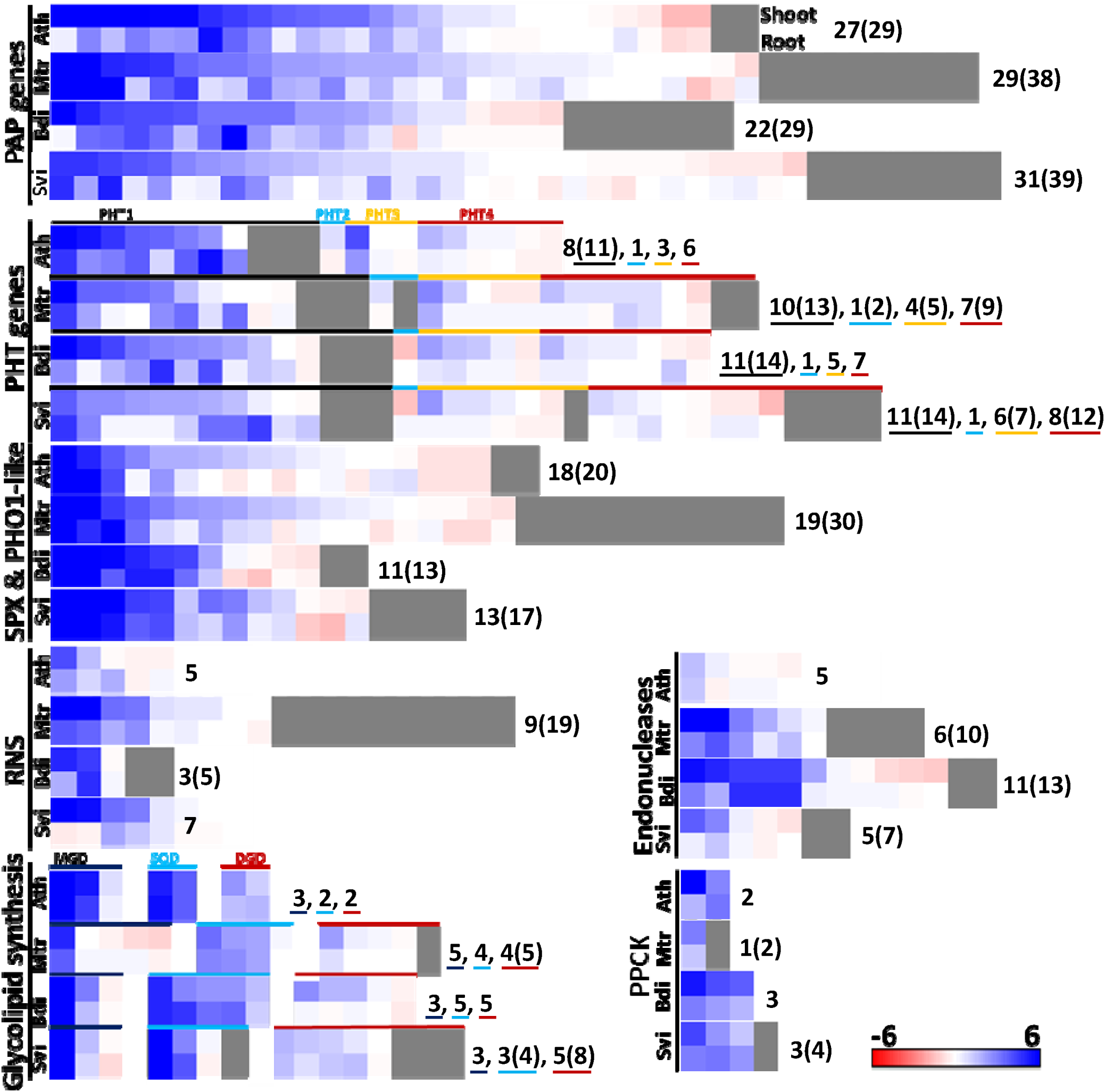
PSI hallmark gene families and their P-limitation response is conserved. *Arabidopsis* (Ath) P-limitation responsive hallmark gene families and their homologous genes in *Medicago* (Mtr), *Brachypodium* (Bdi) and *Setaria* (Si) are shown in a false color scale of log_2_ transformed fold change as blue (≥6) to white (=0) to red (≤-6). Each square represents one gene. The number of each gene family members detected by RNA-Seq is shown and total number of gene family members known including not detected in our RNA-seq are shown in parenthesis. The 1^st^ row for each species represents shoot and 2^nd^ row represents root. Purple acid phosphatases (PAP), phosphate transporters (PHT), SPX domain containing and PHO1-like genes, ribonucleases (RNS), endonucleases, glycolipid synthesis (MGD-monogalactosyl diacylglycerol, SQD- Sulfoquinovosyl diacylglycerol and DGD-Digalactosyl diacylglycerol) and Phosphoenolpyruvate carboxylase kinases (PPCK) genes are shown for all four plant species.

**Figure 5.**
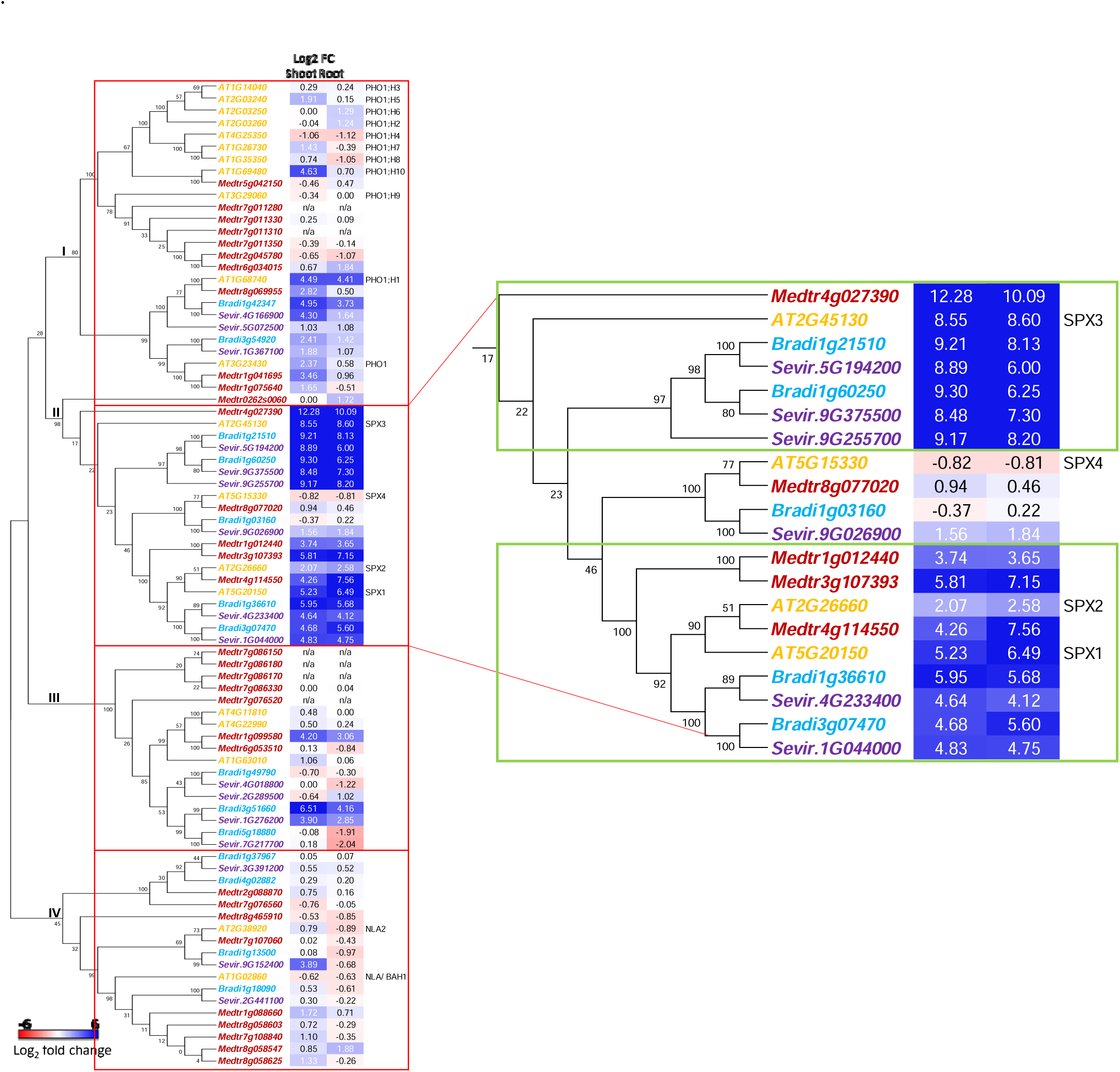
Molecular Phylogenetic analysis of SPX and PHO1-like genes and their response to P-limitation. Maximum likelihood phylogenetic tree was constructed for *Arabidopsis* SPX and PHO1-like genes and their homologs in *Arabidopsis*, *Medicago*, *Brachypodium* and *Setaria*. Protein sequences for all genes were obtained from EnsemblPlants database (Kersey et al., 2016) and the phylogenetic tree was constructed using MEGA6 (Tamura et al., 2013). For each gene, corresponding log_2_ fold change (P-limitation *vs* full-nutrition) values from RNA-seq analysis in shoot and root are shown in a color scale of blue (≥6) to white (0) to red (≤6). Bootstrap test was performed for 1000 replicates and corresponding values are shown for each node. Clades of the tree are numbered I-IV. Gene IDs for different plant species are distinguished by different colors-orange for *Arabidopsis*, red for *Medicago*, blue for *Brachypodium* and purple for *Setaria*.

SPX and PHO1-like genes are involved in signaling P-limitation and Pi transport (Hamburger et al., 2002; Wang et al., 2009; Secco et al., 2012; Wang et al., 2021). Molecular phylogenetic analysis by the maximum likelihood method revealed four distinct clades of SPX-PHO1 like genes (Figure 5. Clade I contains PHO1 family genes, which contain SPX and EXS domains (Wang et al., 2004). The *Arabidopsis* genome contains 11 PHO1 family members comprising PHO1 and 10 additional members, of which five members including *PHO1* and *PHO1;H1* were induced by >2-fold during P-limitation. PHO1 and PHO;H1 from all four plant species formed a distinct subclade on their phylogenetic tree. *PHO1* and *PHO;H1* in *Arabidopsis* and their orthologs in *M.tr.*, *B.di.* and *S.vi.* were all induced by P-limitation. Conservation of *PHO1* orthologs and their conserved P-limitation response suggest and cement their important roles during P limitation, i.e. maintenance of P homeostasis (Liu et al., 2012). Clade II of the phylogenetic tree contains four SPX domain containing genes in *A.th.*, six in *M.tr.*, five in *B.di.* and six in *S.vi.* (Figure 5). *Arabidopsis* SPX1, SPX2 and SPX3, and their orthologs in the other three species were induced by P-limitation revealing the conserved genes and conserved P-limitation response. In contrast, SPX4 of *A.th.*, as well as its ortholog in *B.di.*, showed slight down-regulation during P-limitation but the orthologs were slightly upregulated in *M.tr.* and *S.vi.*. Clade III of the phylogenetic tree contains major facilitator superfamily (MFS) genes harboring SPX and MSF domain, which can function as uniporters, symporters or antiporters, and are involved in transport of various substrates (Secco et al., 2012). This clade contains three, seven, three and four genes from *A.th.*, *M.tr.*, *B.di.* and *S.vi.*, respectively. Out of the 17 genes in this clade, one *A.th.* and two *M.tr.* genes were slightly induced, and one gene in each of *B.di.* and *S.vi.* were more strongly induced by P-limitation. Clade IV includes genes containing SPX and zinc finger (C3HC4-type RING finger) domain proteins. This clade consists of two, nine, four and three genes from *A.th.*, *M.tr.*, *B.di.* and *S.vi.*, respectively. Medtr8g465910 came closer to AT2G38920 in the phylogenetic tree because of the similarity in the RING domain, but is separated as an outgroup because it does not have SPX or EXS domain. Two *B.di.*, one *S.vi.* and two *M.tr.* genes form a distinct subclade in clade IV and contain EXS-superfamily domain but no RING-domain. *At1g02860* (*NLA/BAH1*) is slightly (∼2-fold) down-regulated during P-limitation, most likely because it is the target of P-limitation induced *miR827* (Hsieh et al., 2009; Pant et al., 2009). AT2G38920 (NLA2), a probable E3 ubiquitin-protein ligase, forms a subclade together with its orthologs in *M.tr.*, *B.di.* and *S.vi.* Combining gene expression data with phylogenetic analysis revealed that expressologs (putative functional orthologs) from four different species grouped together forming distinct branches and sub-branches in the phylogenetic tree (Figure 5, S4 and S5).

Similarly, phylogenetic trees were constructed for glycolipid biosynthesis genes (Figure S5). Plants undergo extensive lipid remodeling during phosphorus (P) limitation by replacement of phospholipids with nonphosphorous glycolipids (Hartel et al., 2000; Pant et al., 2015). Expression analysis of these genes during P-limitation revealed that their P-limitation induction is largely conserved across all four plant species (Figure S5 and S6). Phylogenetics combined with gene expression analysis also revealed gene duplications and functional redundancy for many glycolipid synthesis genes (Figure S5A).

### Divergence in P-limitation response of *Arabidopsis* orthologs in other species

During evolution, genes may retain their function, undergo neofunctionalization or subfunctionalization or lose their function. Overview of P-starvation inducible (PSI) genes in *Arabidopsis* and their homologs in four plant species revealed the similarity and diversity of the *PSI* genes and their homologs in different species (Figure 6A). To compare the degree of gene conservation and their P-starvation response among the four plant species, potential orthologs of top 200 *Arabidopsis* PSI genes from both shoot and root (total of 308 unique genes) were identified in *M.tr.*, *B.di.* and *S.vi.*. Out of 308 *A.th. PSI* genes, one or more homologs were found for 251 (81%), 245 (79.5%) and 244 (79.2%) genes in *M.tr.*, *B.di.* and *S.vi.*, respectively. P-limitation induction of these genes was analyzed based on our RNA-seq expression data. Although there were multiple homologs of *A.th. PSI* genes in the other three species, not all the close homologs were P-limitation induced. This comparison of the P-limitation induction of *A.th.* and its close homologs in the other three species revealed that only 133 (53%), 112 (46%) and 121 (50%) genes with identifiable homologs retained their P-limitation induction (≥2-fold cutoff) in *M.tr.*, *B.di.* and *S.vi.*, respectively (Figure 6B). Fifty-six *PSI* genes in *A.th.* had homologs (orthologs identified using PLAZA3.0) in *M.tr., B.di.* and *S.vi.* that were not P-limitation inducible (Figure 6B). The molecular functional categorization of these genes revealed 14 different functional categories (Figure 6C). Most of these genes belonged to other transferase activity (16%), other binding (13%), hydrolase activity (11%), and DNA or RNA binding (10%). There were also gene categories annotated as protein binding, other enzyme activity, unknown molecular function, kinase activity and transcription factor activity among others (Figure 6C). MADS box family transcription factor (*AT5G55690*), transcription elongation factor SPT5 (*AT2g34210*), nuclear factor NF-YA7 (*At1g30500*), transcription factor ATBZIP12 (*AT2G41070*), basic helix-loop-helix (BHLH) DNA binding factor (*At1g27660*) and ethylene response factor ERF/AP2 (*AT1G71130*) are transcription factors which were induced by P-limitation in *A.th.*, but their closest homologs in *M.tr.*, *B. di.* And *S.vi.* were not induced. Also, a plant natriuretic peptide (*PNP-A/ AT2G18660*) was induced in P-limited *Arabidopsis* root but its closest homologs in other three species were not.

**Figure 6.**
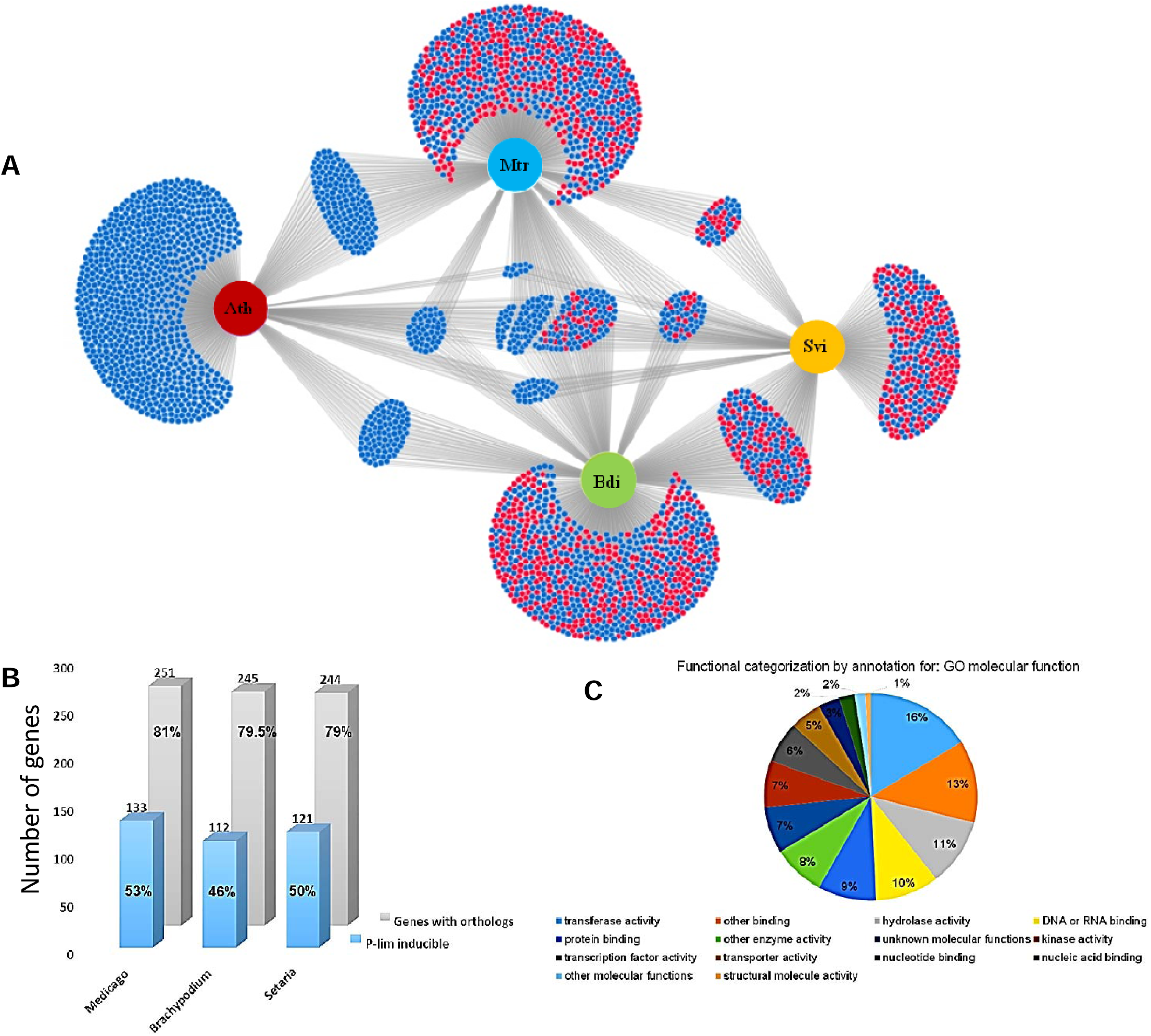
Overview of P-starvation inducible (PSI) genes in *Arabidopsis* and their homologs. (A) Similarity and diversity of *Arabidopsis* PSI gene homologs in other plant species. DiVenn diagram showing *Ath* (*Arabidopsis*) PSI genes (>3 FC) and *Ath* homologs of *Mtr* (*Medicago*), *Bdi* (*Brachypodium*) and *Svi* (*Setaria*) PSI genes in shoot samples. Up (>1 FC) and downregulated (<1 FC) genes are shown as blue and red nodes respectively. Homologs were identified using plant ensemble database. Fold change was carefully selected to include the overall picture of *Arabidopsis* PSI genes, their homologs and expression patterns in other species. (B) Comparison of homologous genes induced by P starvation. Top 200 PSI genes from shoot and root (308 altogether) of *Arabidopsis* and their homologs in *Medicago* (Mt), *Brachypodium* (Bd) and *Setaria* (Sv) are shown. Gray bars show the number of *Arabidopsis* genes for which the homologs are present, while the blue bars show the number of those genes whose homologs were also P starvation induced in *Medicago*, *Brachypodium* and *Setaria*. (C) Gene ontology classification of *Arabidopsis* PSI genes that have close homologs in *Mtr, Bdi* or *Si* but were not P-limitation inducible.

### P-limitation induced but non-conserved transcriptome across four species

Plant genomes contains lineage-specific genes that are not conserved in other species. Lineage-specific genes originate either by *de novo* origination or differential gene loss and retention or from non-lineage-specific paralogs or from lineage-specific genes which get modified from the DNA exapted from transposable elements (Donoghue et al., 2011). Many of the lineage-specific genes identified in *Arabidopsis* were found to be abiotic stress responsive previously (Donoghue et al., 2011). To identify the important species-specific *PSI* genes, top 200 *PSI* genes in each species were selected. The full-length protein or cDNA sequences were taken from TAIR for *Arabidopsis* and Phytozome for *Medicago*, *Brachypodium* and *Setaria*. BLAST searches for each gene were performed with a stringent e-value cutoff of 0.1 in TAIR, Phytozome and NCBI gene bank. Genes that did not have BLAST hits in other three species with e-value cutoff of <0.1 were considered non-conserved genes across the four species. Using this approach, we identified 31, 47, 15 and 36 non-conserved P-limitation induced genes in *A.th.*, *M.tr.*, *B.di.* and *S.vi.*, respectively (Table S2, an excerpt is shown in Figure S7). Though not conserved in the other three species, some of these genes were conserved in their respective taxonomic orders. AT5G17340, a putative membrane lipoprotein, is conserved in the species of order Brassicales-Malvales, suggesting its relatively recent evolutionary origin and function. Most of these P-limitation induced species-specific genes are not well characterized to date. Functional categorization by GO molecular function annotation for *A.th.* revealed that 18 (72%) genes were unknown for molecular function, 4% had protein binding, 12% had other binding and another 12% had other molecular functions. Similarly, 30 (63%) out of 47 genes in *M.tr.* were annotated as hypothetical protein with unknown function, two genes were annotated as leguminosin proline-rich group 669 secreted peptide. *B.di.* and *S.vi.* genes were checked in AgriGO, a GO analysis toolkit (Du et al., 2010), and none of these species-specific genes could be mapped with any gene ontology category, suggesting that the functions of these genes are not known yet.

### Coordinated repression of light reaction-related genes in C3 but not C4 species during P-limitation

Genes involved in the photosynthesis photosystem II (PSII) such as *At1g14150*, *At1g76450*, *At2g28605*, *At2g39470*, *At3g01440*, *At4g21280*, *At4g28660* and *At5g11450* were repressed by P-limitation in Arabidopsis. Similarly, their homologs in *Medicago*, *Brachypodium* and *Setaria* such as *Medtr3g096750*, *Medtr5g041910*, *Medtr1g069465 Medtr3g108040*, *Medtr2g082580*, *Medtr3g108040*, *Bradi5g10500*, *Bradi3g40190*, *Bradi3g39830*, *Bradi2g52730*, *Bradi1g77047*, *Bradi5g16050*, *Bradi4g07440*, *Si011144m.g*, *Si010944m.g*, *Si037067m.g*, *Si020704m.g*, *Si002811m.g* and *Si002383m.g*, were also repressed by P-limitation. Calvin cycle-related genes that are repressed by P-limitation include *At2g47400*, *At5g38420*, *At5g38430*, *At1g56190*, *At3g04790*, *Medtr7g007120*, *Medtr4g093620*, *Medtr2g035660*, *Medtr7g114900*, *Medtr6g018310*, *Medtr6g018300*, *Bradi1g64540*, *Bradi3g27278*, *Bradi3g26391*, *Bradi5g04080*, *Bradi4g36310*, *Bradi4g09120*, *Si038742m.g*, and *Si038812m.g*. Repression of photosynthesis related genes suggests decreased photosynthesis during P-limitation. A set of light reaction related genes were repressed in all three species but not in C4 Setaria (Figure S8) suggesting distinct P starvation response strategies in C4 species.

### Identification of P-limitation core marker genes in evolutionarily distant *Prunus* species

To identify P-limitation core marker genes in *Prunus* species, commonly upregulated genes (>5-fold) in *Ath, M.tr., B.di.* and *S.vi.* were selected. List of *Prunus* homologs (1-to-1 orthologues, 1-to-many orthologues, and many-to-many orthologues of *Arabidopsis* to Prunus, *Medicago* to Prunus, *Brachypodium* to *Prunus* and *Setaria* to *Prunus* orthologs) of *PSI* genes in each species were identified from ensemble database. The common *Prunus* genes were identified from all these four lists and the genes were named based on their best match (homology) to the *Arabidopsis* genes. RT-qPCR based gene expression analysis of selected 14 genes was performed (Figure 7). Of these, P-limitation induction of 93 % *Prunus* genes was verified by RT-qPCR (Figure 7B). These 13 of 14 *Prunus* genes which showed induced expression during P-limitation are most likely the functional orthologs of their *Arabidopsis*, *Medicago*, *Brachypodium* and *Setaria* counterparts (expressologs). Previous studies have also experimentally proven that predicted expressologs are indeed functional orthologs, while nonexpressologs or nonfunctionalized orthologs are not (Das et al., 2016; Vercruysse et al., 2020). Examples of these P-limitation marker genes in *Prunus* include genes encoding for phosphoenolpyruvate carboxylase kinase 1, myb-like transcription factor, Pyridoxal phosphate phosphatase, Major facilitator superfamily protein, EXS (ERD1/XPR1/SYG1) family protein, inorganic pyrophosphatase 1, phosphate transporter, SPX domain protein 3, phospholipase D P2, purple acid phosphatase 16, pyrophosphorylase 4, purple acid phosphatase, and sulfoquinovosyldiacylglycerol 2.

**Figure 7.**
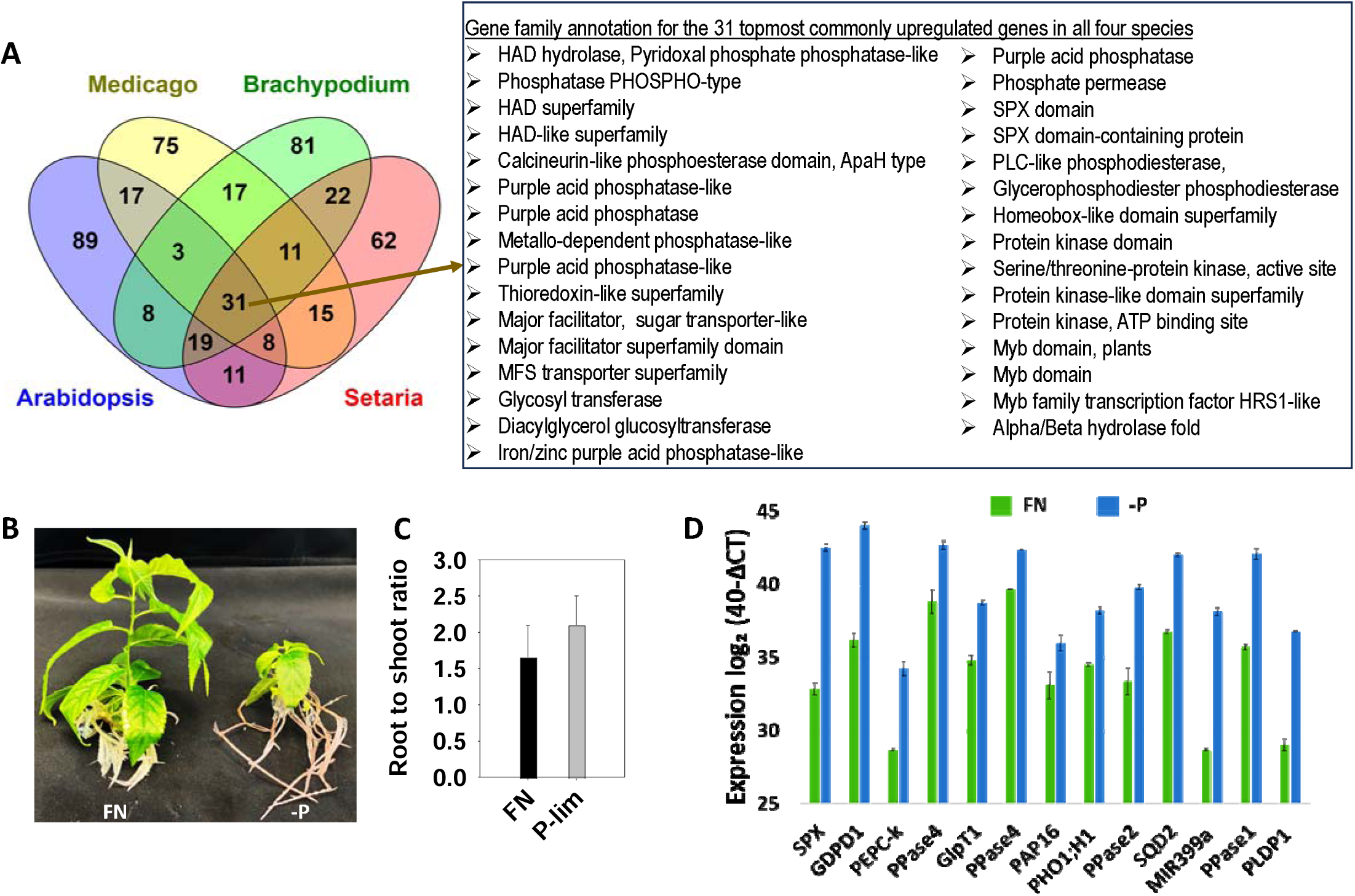
RT-qPCR verification of phosphate starvation inducible genes in *Prunus*. Venn diagram showing the top 100 phosphate starvation inducible (PSI) gene families in the shoots of *Arabidopsis*, *Medicago*, *Brachypodium* and *Setaria*. Gene family was based on InterPro description, where the same gene can be assigned to multiple gene families based on their evolution, related functions, sequence homology or similarities in their structure. Gene families of the 31 topmost commonly upregulated genes in all four plant species are shown on the right. *Prunus* homologs of the commonly upregulated genes in all four plant species were identified and selected for RT-qPCR gene expression analysis. (B) *Prunus* plants grown under full nutrition (FN) and P-limited (-P) condition. (C) Root to shoot ratio of fresh biomass of the plants grown under FN (black bar) and -P (gray bar) before harvesting. (D) RT-qPCR quantification of *Prunus* gene expression in shoots of FN (green) and -P (blue) samples are plotted on a log_2_ scale as 40-ΔC_T_ with ΔC_T_ being the difference between the C_T_ (threshold cycle number) of the gene of interest and the *Ubiquitin* reference gene. Error bars represent standard error of three biological replicates.

## DISCUSSION

Most information about the plant P-limitation response comes from the model plant *Arabidopsis thaliana*, which was used as an anchor model plant species in this study to compare the transcriptomic P-limitation responses in other plant species. Identification of the functional orthologs of well-characterized *Arabidopsis* genes in other plants will help to translate this knowledge from model plant species to crops. As proof of concept, a subset of the core P-limitation responsive genes (expressologs) identified from the four plant species were confirmed for their induced expression in *Prunus* species. Comparable transcriptomic datasets for shoots and roots were generated for the identification of P-status responsive genes across the four different model plant species *Arabidopsis thaliana*, *Medicago truncatula* (model legume), *Brachypodium distachyon* (C_3_ model grass) and *Setaria viridis* (C_4_ model grass) by growing all the plants side-by-side in pots in a growth chamber. Analysis of the RNA-seq expression data revealed conserved and non-conserved transcripts responding to P-limitation. Using our data generated in *Arabidopsis* many novel P-starvation-inducible genes were identified and validated in our recent study (Scheible et al., 2023). Here, we also identified novel *PSI* genes in annotated as well as unannotated genomic regions of *Medicago*, *Brachypodium* and *Setaria*. Induction of the novel *PSI* genes during P-limitation was verified by RT-qPCR analysis and visualizing RNA-seq reads in the integrative genomics viewer (IGV). Visualization of the RNA-seq data in IGV also revealed that many genes undergo alternative splicing during P-limitation in all four species. This is consistent with the previous findings that several gene transcripts were alternately spliced out during P-limitation in *Arabidopsis* (Scheible et al., 2023). Alternative splicing events during P-limitation was also confirmed by PCR and RT-qPCR analysis (Scheible et al., 2023). Alternative splicing is known to diversify the transcriptome and proteome and plays vital role in maintaining nutrient homeostasis in plants (Dong et al., 2018; Jeon et al., 2022).

Most of the known hallmark genes of P-starvation response in *Arabidopsis* and their orthologs were found to be induced during P-limitation in all four plant species. Majority of the highly P-starvation inducible genes were attenuated after P resupply to the P-starved plants confirming their regulation directly by P availability, rather than indirectly by downstream events including altered metabolism, growth, and development. Gene families that play an important role in phosphate signaling and transport, and those involved in conserved and species-specific biological pathways were studied in detail using molecular phylogenetic analysis.

Here we also report the establishment of experimental protocol to study the P-limitation responsive genes in *Prunus* using tissue culture plants. RT-qPCR verification of *PSI* core genes identified by this study in *Prunus* also showed the usefulness of this comparative transcriptomics study in pinpointing the functional orthologs in additional plant species such as the evolutionarily distant *Prunus*. Because of the gene or genome duplications events in evolutionarily distant species it is difficult to identify the functional orthologs among one-to-many or many-to-many orthologs. Since expression pattern similarities support the prediction of orthologs retaining common functions after gene duplication, identification of expressologs is key to pinpoint the functional orthologs in different species (Das et al., 2016; Vercruysse et al., 2020). Identification of *Prunus* expressologs during P-limitation and their verification by RT-qPCR reveals the usefulness of this comparative transcriptomics study in translating knowledge to other distantly related plant species. Conserved expression of these *PSI* genes in diverse plant species reveals their essential molecular function to maintain P-homeostasis in plants. In the same manner, this method of identification of functional orthologs (expressologs) can be used to pinpoint, for example, any biotic or abiotic stress-responsive genes in a wide range of species.

Although the general P-limitation response of many *PSI* genes was similar in all four plant species, the number of genes in each family and the extent of their P-response was different in each species. This could be due to unique gene duplication events in the lineage of each species, and non-, neo- or sub-functionalization of the gene family members after the structural and functional divergence during genome evolution (Teufel et al., 2016). We identified many *Arabidopsis* P-starvation inducible genes whose homologs in the other three species were not induced by P-limitation. We also identified many species-specific P starvation-inducible genes, which do not have close homologs in the other three plant species. Most of these genes are poorly annotated and their biological function is still unknown. Very high induction of these species-specific genes by P-limitation indicates that there exist some species-specific mechanisms for P acquisition and homeostasis in different plants.

P-limitation led to coordinated changes in the expression of genes involved in P uptake and internal P mobilization in all four species. Most highly P starvation-inducible genes in all four species include many hallmark genes of P-starvation response such as *SPX* genes that are involved in P-signaling, phosphate transporters, purple acid phosphatases, phospholipase and glycolipid synthases, and transcriptional regulators. A previous study in *Arabidopsis* showed that many *PSI* genes involved in above mentioned pathways, phospholipase genes for example, were reversibly induced by P-limitation (Morcuende et al., 2007). Phylogenetic analysis combined with the gene expression during P-limitation helped to identify functional orthologs (expressologs) with similar expression patterns in other species. Phylogenetic analysis of the SPX domain proteins revealed that NLAs formed a separate branch because it contained both SPX and RING-domains. Downregulation of *AtNLA* and its orthologs during P-limitation suggested its conserved function across the diverse plant species. NLA is known to regulate Pi transport by mediating posttranscriptional degradation of PHT1s (Kant et al., 2011; Lin et al., 2013; Bucher and Fabianska, 2016). Many pathogenesis-related (PR) genes were induced during P-limitation in all four species represented in our dataset. PR genes belong to 17 different families (PR-1 to -17) of PR proteins identified from various flowering plants (Liu and Ekramoddoullah, 2006). Based on these results and additional experimental findings, we were able to discover that P-limited plants are more resistant to pathogens and drought stress, which is mediated by PHR1–PHL1 in *Arabidopsis* (Scheible et al., 2023).

Genes involved in essential metabolic pathways in plants such as photosynthesis, lipid metabolism and secondary metabolism were found to be regulated by P-limitation in all four plant species. In *Arabidopsis*, phosphate starvation repressed transcripts involved in photosynthesis were not reverted after P resupply suggesting that these changes are indirect responses and may be linked to lower demand for photosynthate and increased sugar levels during P-limitation (Morcuende et al., 2007). Photorespiration-related genes were induced during P-limitation. This is consistent with the increased photorespiration during P-limitation (Pant et al., 2015). Gene ontology (GO) analysis of the top *PSI* genes in shoot and root of four plant species revealed genes associated with diverse biological processes including lipid metabolism, signaling, cell communication, cellular response to phosphate starvation and nutrient levels, phosphate transport, negative regulation of transcription, nucleoside, nucleotide and nucleic acid metabolic process, nitrogen compound metabolic process, and other metabolic process. This shows alteration in various biological processes to maximize the P uptake and maintain P homeostasis during P-limitation in plants. However, many PRGTs could not be assigned to any GO functional annotation because their functional domain is not characterized yet. Most of these genes were highly induced during P-limitation but not expressed during P-sufficient condition. While detailed functional characterization of these different and unique novel set of genes expressed during P-limitation is not within the scope of this study, further work on their functional characterization could shed light on the currently unknown molecular mechanisms and biological processes. Novel findings from this study will enrich our knowledge on how different plants adapt to P-limitation and can be used in translational research to engineer more P-efficient crop species.

## CONCLUSION

Comparable transcriptomics datasets were generated for the identification of sets of phosphorus (P)-status responsive genes in four model plant species as *Arabidopsis thaliana* (Brassicaceae), *Medicago truncatula* (legume), *Brachypodium distachyon* (C3 grass, like rice or wheat) *and Setaria viridis* (C4 grass, like switchgrass or maize), including potential regulators of P-responses that are conserved across the species. Genome-wide transcriptional responses in shoots and roots of the four species grown under P-sufficient and P-limitation conditions using RNA sequencing revealed conserved and non-conserved P-limitation responsive genes, their conserved and non-conserved responsiveness to P-limitation, as well as non-conserved unique genes that respond to P-limitation. The use of RNA-seq identified novel annotated and unannotated P-responsive genes that were not identified by previous studies. A subset of *Medicago* and *Brachypodium PSI* genes identified by RNA-seq were analyzed using RT-qPCR, confirming their P starvation responses and reversed expression after P resupply to the P-starved plants. These RT-qPCR verified genes that were found to be directly regulated by P availability in independent experiments can serve as important P-limitation marker genes in legumes and grasses. Phylogenetics, combined with gene expression analysis, revealed the conserved and lineage or species-specific responses to P-limitation. This study provides proof-of-concept that this comparative transcriptomics study can be successfully used to pinpoint the functional orthologs (expressologs) in diverse plant species including the *Prunus* (tree) species. P-limitation induced hallmark genes and primary metabolism-related genes such as sucrose and starch biosynthesis, lipid metabolism, P-salvage related genes, and their P-limitation response is largely conserved, though the number of genes belonging to each family and their extent of P-limitation response was different. Although many genes and processes are conserved across the species, there exists considerable species-specific response to P-limitation. These findings will be helpful to translate knowledge from model organisms to other crop species and engineering of the P-efficient crops.

### SUPPLEMENTARY DATA

Supplemental Figure S1. Examples of alternatively spliced genes during P-limitation.

Supplemental Figure S2. Induction of PSI genes reversed after P-resupply to P-starved plants in *Brachypodium*.

Supplemental Figure S3. Overview of transcriptional regulator (TR) genes responding to phosphorus limitation in four plant species.

Supplemental Figure S4. Molecular phylogenetic analysis of *PHT1* and *PHT3* genes, and their response to P-limitation.

Supplemental Figure S5. Molecular phylogenetic analysis of glycolipid synthesis genes.

Supplemental Figure S6. P starvation-response of lipid metabolism-related genes in four plant species

Supplemental Figure S7. P-limitation induced non-conserved genes from *Arabidopsis, Medicago, Brachypodium* and *Setaria*.

Supplemental Figure S8. P-limitation response of photosynthesis (light reactions) genes.

Supplemental Table S1. Output file from the RNA-seq analysis, phosphate responsive gene transcripts in all four species and primers used in the present study.

Supplemental Table S2. Non-conserved P-limitation induced genes across the four species.

## ACKNOWLEDGEMENTS

We thank Yuhong Tang and Stacy Allen for their technical assistance during RNA-seq analysis and Sylvia Warner for technical assistance in the laboratory.

## AUTHOR CONTRIBUTIONS

WRS: conceptualization for RNA-seq; PP and HD: conceptualization for remaining part; PP and WRS: methodology; NK: processed RNA-seq raw data; WRS, and PP: formal analysis, data curation and figures; WRS and PP: investigation; WRS, and HD: resources; WRS and PP: writing - original draft; WRS, PP, and HD: writing, review, and editing; WRS and HD: funding acquisition.

## CONFLICT OF INTEREST

The authors declare no conflict of interest.

## FUNDING

This work was supported by the United States Department of Agriculture, Agricultural Research Service, Floral and Nursery Plants Research Unit of National Arboretum, USA, and Noble Foundation, Oklahoma, USA.

## DATA AVAILABILITY

All the raw sequencing data generated in this study are deposited in the National Center for Biotechnology Information Sequence Read Archive Database under BioProject ID: PRJNA1076492.

## Notes

### Competing Interest Statement

The authors have declared no competing interest.

